# MIND: Multimodal Integration with Neighbourhood-aware Distributions

**DOI:** 10.1101/2025.09.15.676314

**Authors:** Hanwen Xing, Christopher Yau

## Abstract

Multi-omics profiling has become a powerful tool for biomedical applications such as cancer patient stratification and clustering. However, the characterisation and integration of multi-omics data remain challenging because of missingness and inherent heterogeneity. Methods such as imputation and sample exclusion often rely on strong assumptions that could potentially lead to information loss or distortion. To address these limitations, we propose a multi-omics integration framework that learns patient-specific embeddings from incomplete multiomics data based on a multimodal Variational Autoencoder with a data-driven prior. Specifically, we inject neighbourhood structure of the observed dataset encoded as affinity matrices into the prior of embeddings through exponential tilting, and use this prior to penalise the configuration of the latent embeddings based on the discrepancy between the neighbourhood structures in the data spaces and in the latent space. Our proposed method handles high missing rate and unbalanced missingness pattern well, and is robust in the presence of data with a low signal-to-noise ratio. Compared with existing data integration methods, the proposed method achieves better performance on a range of supervised and unsupervised downstreaming tasks on both synthetic and real data.

## 1 Introduction

Multi-omics data integration has become essential for understanding complex biological systems and advancing precision medicine (Shin et al., 2017; Subramanian et al., 2020; Steyaert et al., 2023). Modern high-throughput technologies enable simultaneous profiling of genomics, transcriptomics, proteomics, and epigenomics from the same samples, each providing complementary views of cellular function (Cao et al., 2018; Regner et al., 2021; Fu et al., 2024). However, integrating these heterogeneous, high-dimensional datasets presents significant computational challenges, particularly when dealing with the pervasive missing data patterns characteristic of real-world multi-omics studies.

Existing integration methods face fundamental limitations in handling incomplete data. Network-based approaches often require overlapping observations or rely on ad hoc imputation or graph fusion strategies (Rappoport and Shamir, 2019; Xu et al., 2021; Ma et al., 2025). Matrix factorisation methods assume linear relationships and struggle with complex missing patterns (Shen et al., 2012; Yang and Michailidis, 2016). Although variational autoencoders (VAE) can naturally accommodate missing data (Gayoso et al., 2021), aggregating information from multimodal data with missing values using VAE requires careful design of both the training scheme (Wu and Goodman, 2018) and the modelling architecture (Ballard et al., 2025; Beaude et al., 2025), which can be computationally intensive. Furthermore, unlike network-based approaches, current VAE models do not exploit the neighbourhood structures within multi-omics datasets explicitly. As a result, they may not be able to preserve the intrinsic clustering structure present in biological data.

We present Multimodal Integration with Neighbourhood-aware Distribution (MIND), a lightweight multimodal VAE, to address these limitations: MIND utilises t-SNE-derived affinity matrices to construct a data-driven prior that preserves neighbourhood structures from individual omics modalities, ensuring biologically meaningful patient-level representations while maintaining VAE’s probabilistic modelling framework. Additionally, MIND employs cross-modal regular-isation to encourages consistency and information sharing between modality-specific encodings from the same patient, thereby stabilising the aggregation step while maintaining flexibility for arbitrary missing patterns. We demonstrate MIND’s superior performance across synthetic datasets and real-world applications including The Cancer Genome Atlas (Weinstein et al., 2013), the Childhood Cancer Multi-omics Atlas (Sun et al., 2023), and the Cancer Cell Line Encyclopedia (Ghandi et al., 2019). Our method consistently outperforms existing approaches such as JASMINE (Ballard et al., 2025), IntegrAO (Ma et al., 2025), and MSNE (Xu et al., 2021) in survival prediction, cancer classification, clustering, and data reconstruction tasks, providing a robust framework for multi-omics integration in diverse biological contexts.

## 2 Results

### 2.1 Methods Overview

Our proposed method uses a multimodal VAE architecture to extract patient-level embeddings from incomplete multiomics datasets. These patient-level embeddings are learnt by first mapping all entries in the incomplete dataset to individual embeddings using a collection of modality-specific encoders, and then aggregating available individual embeddings associated with the same patient into a single embedding. Each of the patient-level embeddings is then passed to each of the modality-specific decoders to reconstruct the multiomics observed data, or predict the missing ones. A schematic illustration of the modelling architecture can be found in Fig 1. To align the learnt penitent-level embeddings with the intrinsic clustering structure present in biological data, we put a data-driven prior that preserves neighbourhood structures from individual modalities using t-SNE-derived affinity matrices on the penitent-level embeddings. This prior distribution penalises the configuration of the latent embeddings based on the discrepancy between patients’ neighbourhood structures in the data spaces and the latent space, and favours latent embeddings whose clustering structure resembles a consensus of clustering structures present in individual modalities. Additionally, a cross-modality regularisation that encourages consistency and information-sharing between individual embeddings of the same patient is used to capture inter-modality dependencies of the incomplete multiomics datasets. Implementation details of MIND can be found in Sec 4

**Figure 1:**
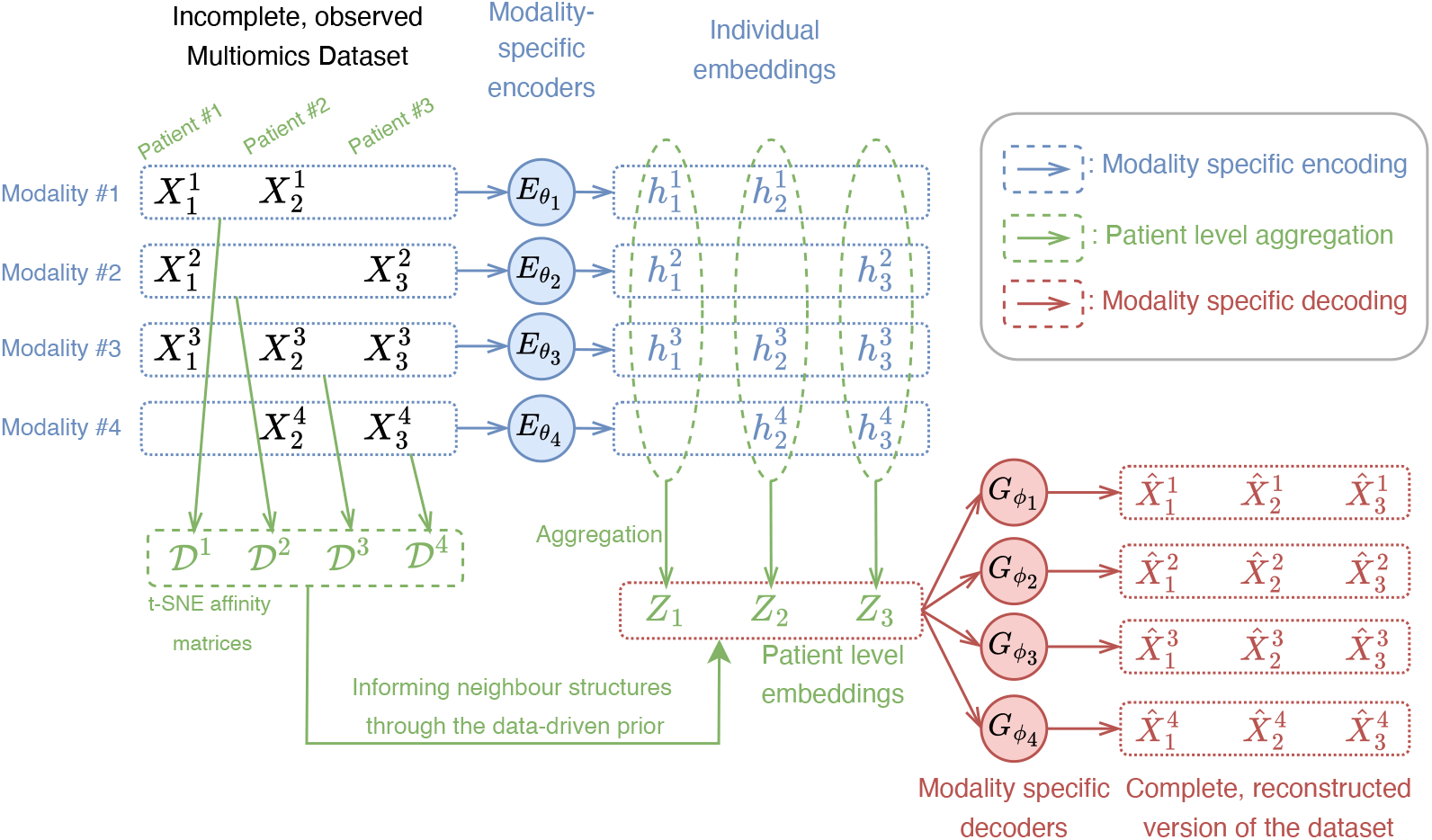
Schematic illustration of MIND. We use an incomplete multiomics dataset with *M* = 4 modalities and *N* = 3 patients as an example. Available 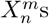 in the incomplete dataset are first mapped to individual embeddings 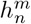s by modality-specific encoders 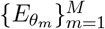 Individual embeddings 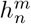 associated with the same patient *n* are then aggregated into a patient level embedding *Z*_*n*_. Each 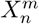 is reconstructed (if available in the incomplete dataset) or predicted (if missing) by passing the patient level embedding *Z*_*n*_ to the modality-specific decoder 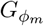 For each modality, a t-SNE affinity matrix 𝒟^*m*^ encoding neighbour structure of the relevant patients is computed. These matrices are used to guide the neighbour structure of patient level *Z*_*n*_s through the prior on *Z*_*n*_s.

### 2.2 Synthetic data

We first demonstrate the efficacy of the proposed method using synthetic multi-omics datasets generated by the `InterSim`CRAN package (Chalise et al., 2016). The synthetic data set consists of three modalities: DNA methylation, mRNA expression, and protein expression. Similarly to Ma et al. (2025), we set the number of patients *N* = 500 and the number of groups *C* = 15. To demonstrate the robustness of the proposed method, we consider two scenarios in which (1) all modalities show clear and consistent clustering patterns and (2) some modalities are more noisy and less informative. We refer to the two scenarios as the low noise regime and the high noise regime, and the corresponding synthetic data sets are generated by passing different parameters to `InterSim.` (See Supp 4.4). Visualization of the two synthetic datasets can be found in Figs 2. To mimic missingness patterns in real data, we follow Ma et al. (2025) and mask subsets of data from each of the modalities as follows: Given an overlap ratio *α*, we first randomly select ⌊*αN* ⌋ patients as common samples (i.e., keep their data intact), then evenly distribute the remaining ⌈(1 − *α*)*N* ⌉ patients among the three modalities as unique samples. For example, when *α* = 0.7 and *N* = 500, each of the three modalities would have 50 unique patients, and the remaining 350 patients are present in all three modalities. We tested our proposed method at overlap ratios *α* ∈ {0.1, 0.2, …, 0.9} in both low and high noise regimes.

**Figure 2:**
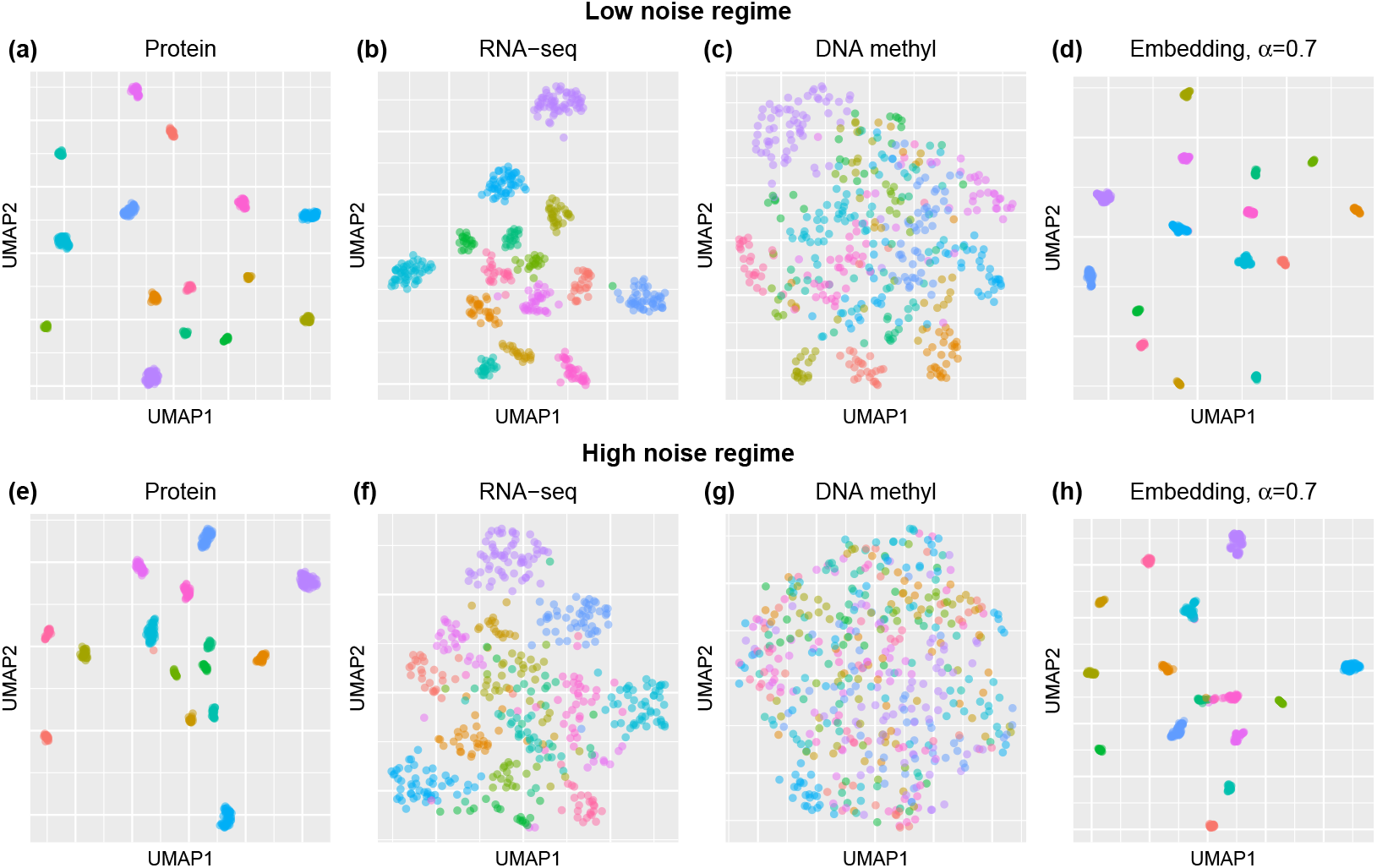
Visualisation of synthetic datasets and embeddings. Top row: Low noise regime. (a-c) UMAP visualisation of the three modalities. (d). UMAP visualisation of the embeddings estimated by MIND at overlap ratio *α* = 0.7. Botton row: Similar figures under the high noise regime.

We compare our proposed method with two current state-of-the-art models, VAE-based JASMINE (Ballard et al., 2025) and network-based IntegraO (Ma et al., 2025). We also include MSNE (Xu et al., 2021) as a baseline. We set the dimension of embeddings *d*_*l*_ = 64 for all methods. We apply these methods to high- and low-noise synthetic datasets and compare the quality of the generated embeddings by investigating their performance on supervised and unsupervised downstreaming tasks. We first investigate the clustering performance of different methods. For each method, we apply spectral clustering with the true number of clusters *C* = 15 to the generated embeddings and compare their clustering results with the true group membership using normalized mutual information (NMI) as a metric. The results at different overlap ratios are reported in Fig 3(a, d). We then consider two supervised tasks, classification and reconstruction. For classification, we train classifiers that predict patient groups from embeddings generated by each method using XGBoost (Chen and Guestrin, 2016), and report 10-fold CV accuracies at different overlap ratios in Fig 3(b, e). For reconstruction, we used the generated embeddings to reconstruct the mask portion of the synthetic data. For each modality, we compute the Pearson correlations between the reconstructed and observed data. We report the averaged correlations of the three modalities in Fig 3(c, f). Here we can only compare these results with those of the VAE-based JASMINE, since network-based IntegrAO and MSNE do not offer this feature. Scatter plots of predicted vs observations given by MIND at *α* = 0.7 are reported in Fig 5.

**Figure 3:**
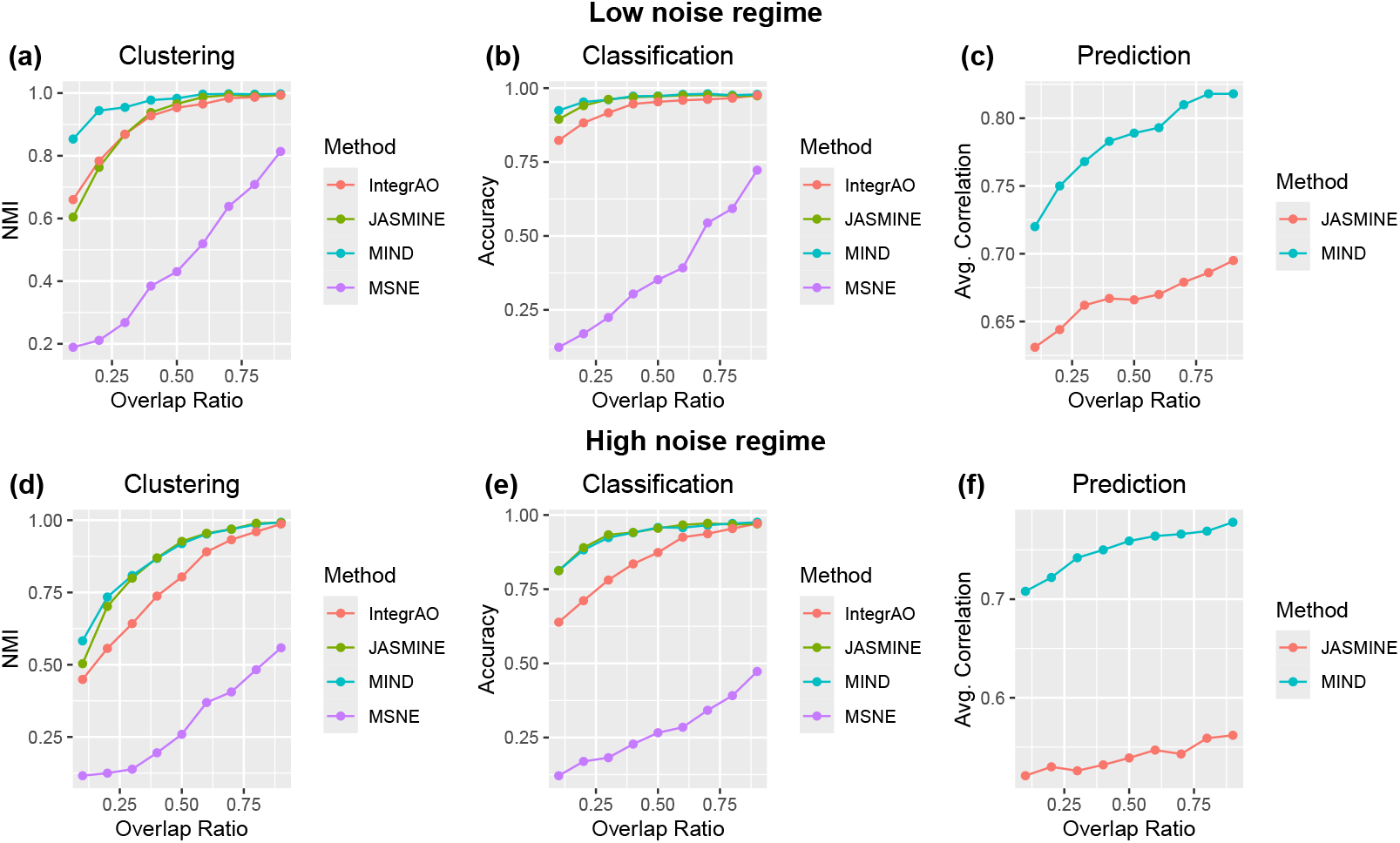
Performance on the three downstream tasks. Top row: Results from the low noise datasets. (a) Normalised Mutual Information vs overlapping data ratio of different methods. (b) Classification accuracy estimated by 10-fold CV of different methods. (c) Prediction accuracy measured by the Pearson correlations between the masked portion of the synthetic data and the predicted, averaged over all modalities. Bottom row: Similar results from the high noise datasets.

**Figure 4:**
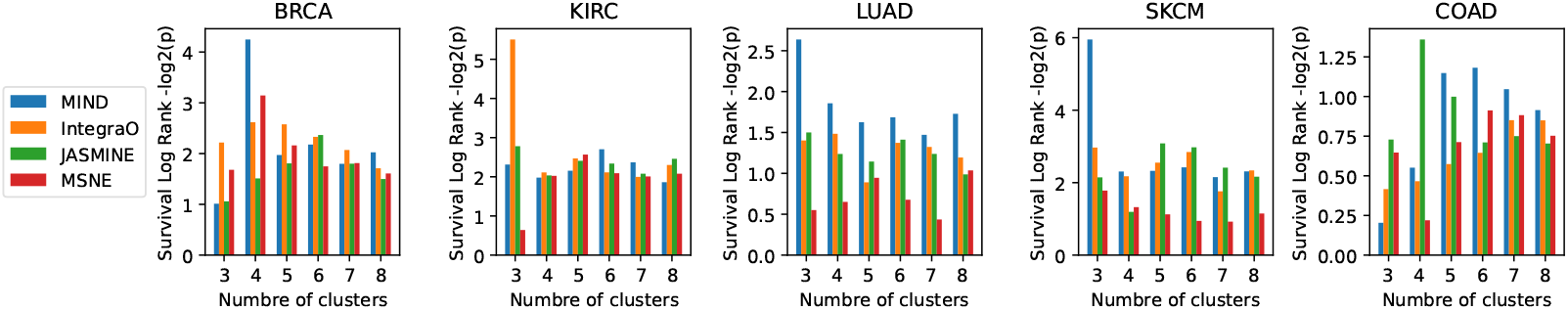
Clustering. Log-rank test for differential survival of among 3-8 clusters of five cancer types in TCGA: BRCA, KIRC, LUAD, SKCM and COAD. Cluster assignments are determined by K-mean. For each method and number of clusters, we report the averaged − log_2_(*p*) value over five replicates, where *p* is the p-value of the pairwise log-rank test.

**Figure 5:**
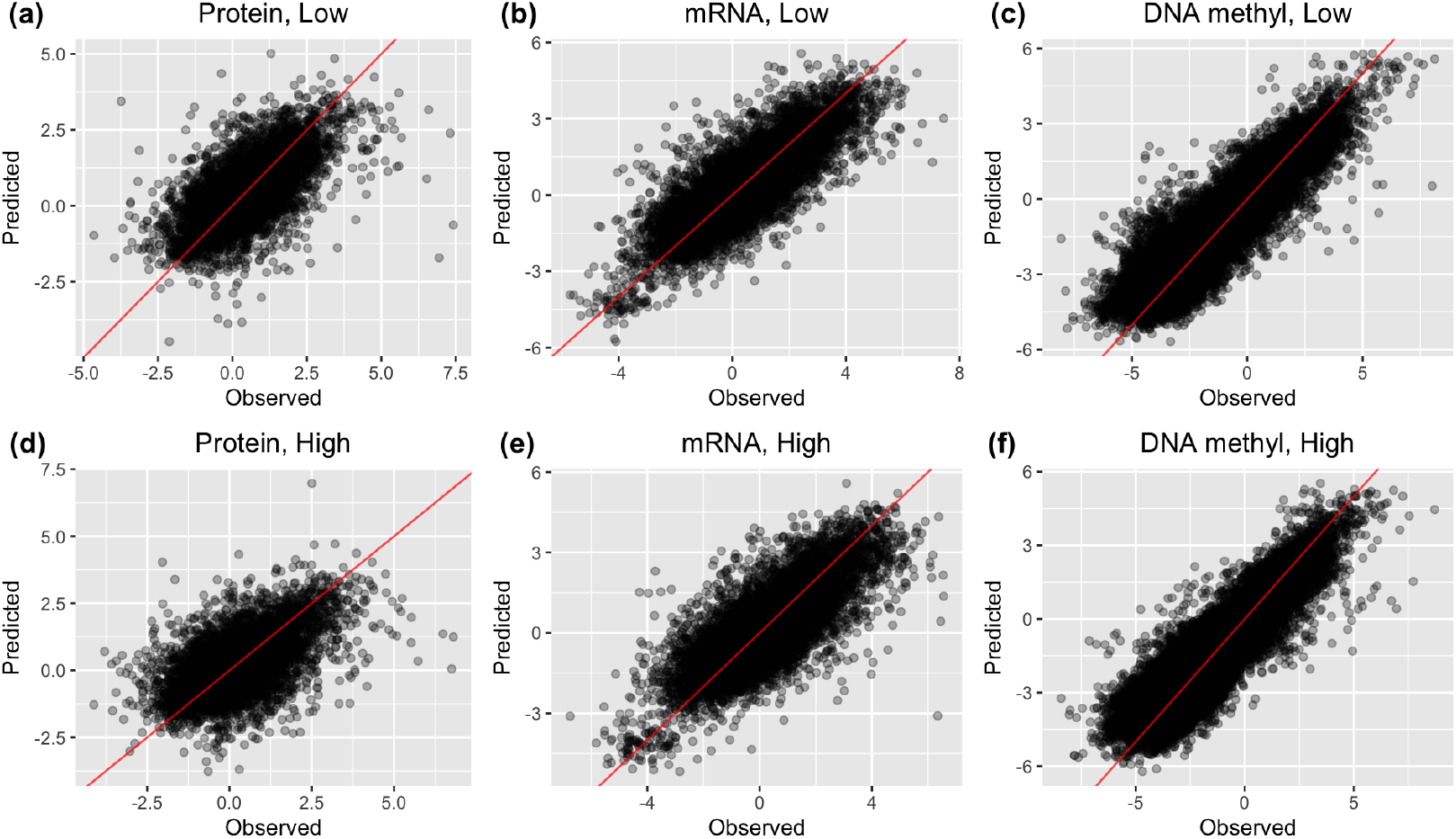
Scatter plots of the predicted value given by MIND vs the observed masked during the training phase at overlap ratio *α* = 0.7. Top: Low noise regime. Bottom: High noise regime.

### 2.3 Pan-cancer prediction tasks

We applied our proposed method to mRNA-seq, miRNA-seq, DNA methylation, copy number variation (CNV) and functional proteomics reverse phase protein array (RPPA) data from The Cancer Genome Atlas (TCGA) (Weinstein et al., 2013). We investigated 17 data sets from TCGA, which show different degrees of missingness. The sizes of the multi-omics datasets and details on data pre-processing can be found in Supp 4.4. In this example, we fix the dimension of latent embedding of our proposed method, IngegrAO, and MSNE to 64. Since the length of JASMINE’s output embeddings has to be a multiple of *M* + 1 where *M* is the total number of modalities, we set its dimension to 66 for a fair comparison. Similarly to the synthetic example, we compare the proposed method with IntegrAO, JASMINE and MSNE on four tasks: survival prediction, cancer stage classification, clustering, and multiomics data reconstruction.

We first investigate the individual datasets. For the survival prediction task, we fit a Coxnet survival model (Simon et al., 2011) using both the embeddings generated by each model and patients’ age and gender as feature vectors to predict the survival status of the patients for each dataset. We use the C-index (Harrell Jr et al., 1996) as a performance metric. For each method and dataset, the C-index calculated using 5-fold CV is reported in Table 1.

**Table 1:**
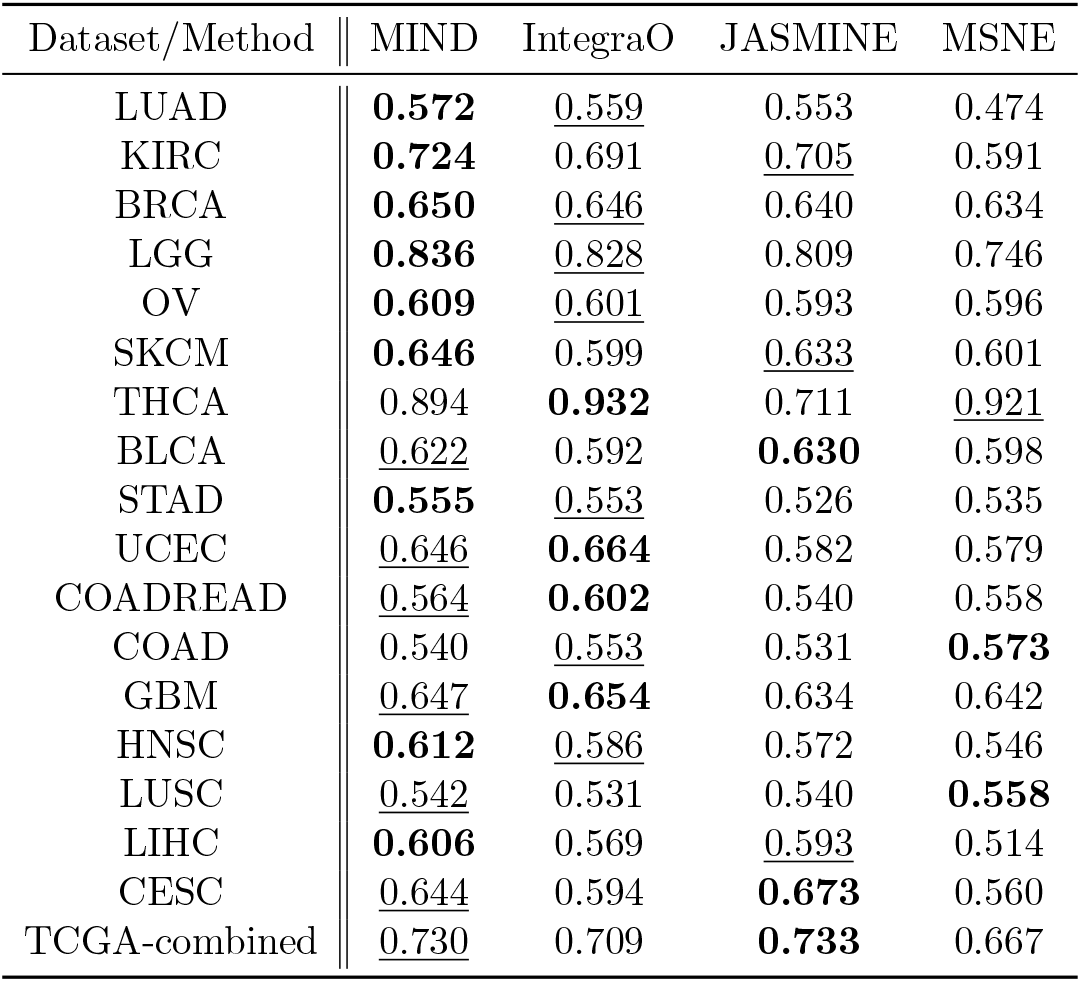
Survival prediction. C-index estimated using 5-fold CV. For this task, penalised Cox’s proportional hazard’s models are trained to predict survival status of patients using patient information (age, gender) and embeddings given by different methods. For each cancer type, the best result is highlighted in **boldface**. Second best is <u>underlined</u>.

| Dataset/Method | MIND | IntegraO | JASMINE | MSNE |
| --- | --- | --- | --- | --- |
| LUAD | **0.572** | <u>0.559</u> | 0.553 | 0.474 |
| KIRC | **0.724** | 0.691 | <u>0.705</u> | 0.591 |
| BRCA | **0.650** | <u>0.646</u> | 0.640 | 0.634 |
| LGG | **0.836** | <u>0.828</u> | 0.809 | 0.746 |
| OV | **0.609** | <u>0.601</u> | 0.593 | 0.596 |
| SKCM | **0.646** | 0.599 | <u>0.633</u> | 0.601 |
| THCA | 0.894 | **0.932** | 0.711 | <u>0.921</u> |
| BLCA | <u>0.622</u> | 0.592 | **0.630** | 0.598 |
| STAD | **0.555** | <u>0.553</u> | 0.526 | 0.535 |
| UCEC | <u>0.646</u> | **0.664** | 0.582 | 0.579 |
| COADREAD | <u>0.564</u> | **0.602** | 0.540 | 0.558 |
| COAD | 0.540 | <u>0.553</u> | 0.531 | **0.573** |
| GBM | <u>0.647</u> | **0.654** | 0.634 | 0.642 |
| HNSC | **0.612** | <u>0.586</u> | 0.572 | 0.546 |
| LUSC | <u>0.542</u> | 0.531 | 0.540 | **0.558** |
| LIHC | **0.606** | 0.569 | <u>0.593</u> | 0.514 |
| CESC | <u>0.644</u> | 0.594 | **0.673** | 0.560 |
| TCGA-combined | <u>0.730</u> | 0.709 | **0.733** | 0.667 |

Similarly, for the cancer stage task, we train XGboost classifiers (Chen and Guestrin, 2016) using the embeddings generated by each model to predict the cancer stage (I-IV) of the patients. We report the 5-fold CV classification accuracy for each method and dataset in Table 2.

**Table 2:** Cancer stage classification. Classification accuracy estimated using 5-fold CV. For this task, XGboost classifiers are trained to predict cancer stages (I-IV) using embeddings given by different methods. For each cancer type, the best result is highlighted in **boldface**. Second best is <u>underlined</u>. - indicates cancer stage information unavailable.

| Dataset/Method | MIND | IntegraO | JASMINE | MSNE |
| --- | --- | --- | --- | --- |
| LUAD | **0.517** | 0.476 | <u>0.505</u> | 0.474 |
| KIRC | <u>0.513</u> | **0.552** | 0.510 | 0.460 |
| BRCA | 0.516 | 0.506 | <u>0.540</u> | **0.545** |
| LGG | - | - | - | - |
| OV | - | - | - | - |
| SKCM | <u>0.456</u> | 0.443 | **0.491** | 0.438 |
| THCA | <u>0.543</u> | 0.540 | **0.573** | 0.533 |
| BLCA | **0.451** | 0.417 | <u>0.432</u> | 0.337 |
| STAD | <u>0.383</u> | 0.377 | **0.425** | 0.376 |
| UCEC | - | - | - | - |
| COADREAD | **0.333** | 0.282 | <u>0.331</u> | 0.324 |
| COAD | 0.334 | 0.324 | <u>0.351</u> | **0.358** |
| GBM | - | - | - | - |
| HNSC | **0.541** | 0.496 | <u>0.529</u> | 0.501 |
| LUSC | **0.472** | 0.442 | <u>0.470</u> | 0.430 |
| LIHC | **0.480** | <u>0.473</u> | 0.472 | 0.371 |
| CESC | <u>0.453</u> | 0.427 | **0.503** | 0.447 |

We then consider the unsupervised clustering task. For each method and each dataset, we apply *k*-means clustering algorithm with cluster number ranging from 3 to 8 to the resulting embeddings. We group patients according to the cluster membership given by *k*-mean, then perform pairwise log rank test based on the estimated membership and the survival status of patients. We use − log_2_(*p*), where *p* is the *p* value of the pairwise log rank test, as a performance metric, since a lower *p* (i.e. higher − log_2_(*p*)) indicates stronger evidence of differences in survival behaviour, and therefore greater inter-group differences. The results of the five datasets BRCA, KIRC, LUAD, SKCM and COAD investigated in Ballard et al. (2025) are reported in Fig. 4. The complete results can be found in Fig 6.

**Figure 6:**
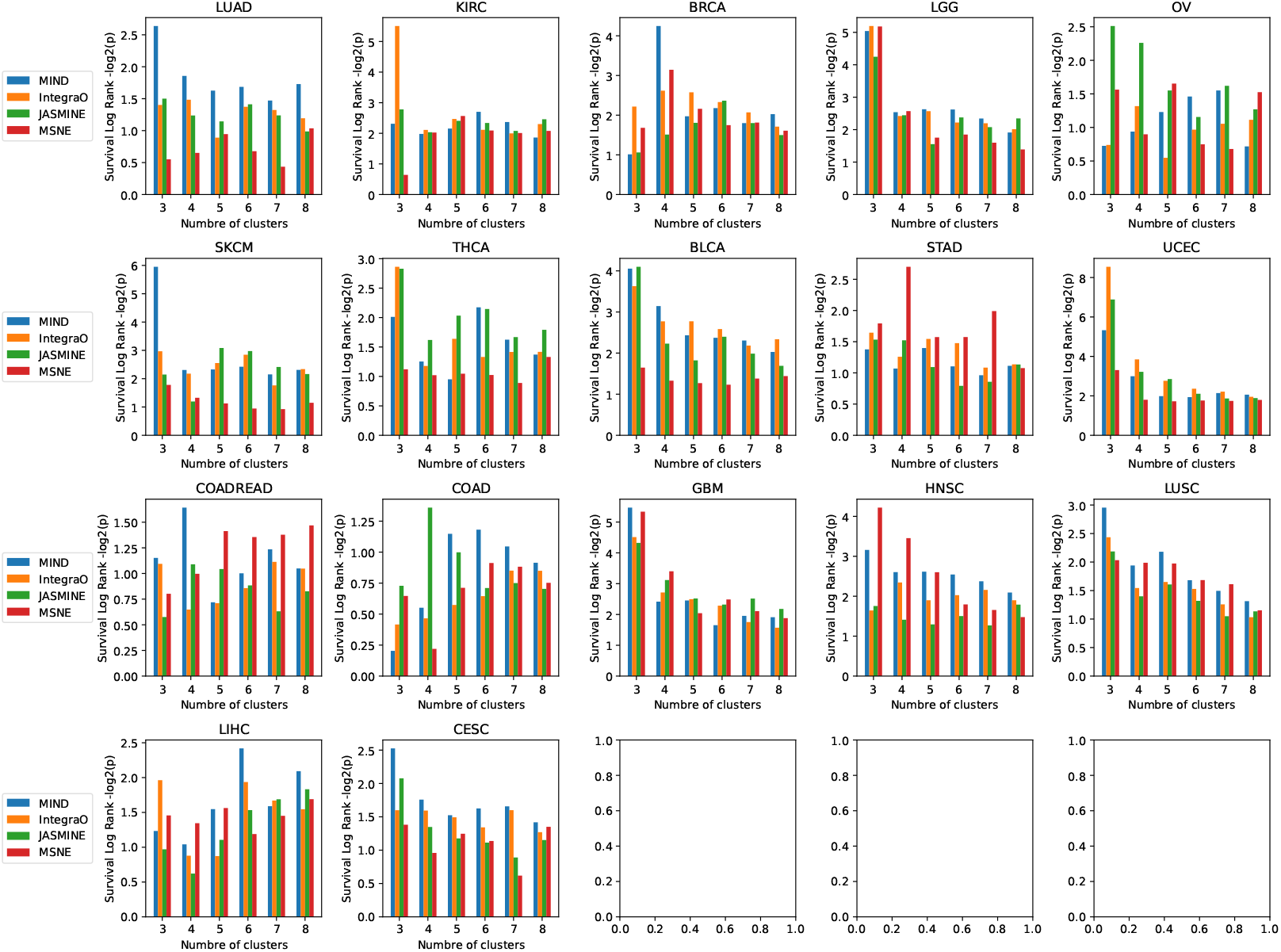
Log-rank test for differential survival of the 17 cancer types among 3-10 clusters. Cluster assignments are determined by K-mean. For each method and number of clusters, we report the averaged − log_2_(*p*) value of the pairwise log-rank tests.

Finally, we compare prediction performance between our proposed method and JASMINE. Recall that IntegraO and MSNE do not have this feature. Here, for each modality of each dataset, we first randomly mask 10% of its data subject to the constraint that, for every dataset, each patient must be present in at least one modality of the resulting masked multiomics dataset. We then train both our proposed method and JASMINE using the masked datasets, predict the masked data using the learned embeddings, and compare the predictions with the observed values. Since the final embedding of JASMINE is obtained by concatenating *M* modality-specific embeddings and a global embedding, a embedding vector of length 66 means that each of its modality-specific embedding only has length 11, which could be too stringent to accurately reconstruct data from each modality. We therefore additionally fit an augmented JASMINE such that each modality-specific embedding has length 64 (i.e. the final embedding has length 384). For each modality of each dataset, we computed the Pearson correlation between the predicted and observed data. We report the averaged Pearson correlation of all five modalities in Table 3.

**Table 3:** Reconstruction. Pearson correlations between prediction and observed values averaged over all five modalities. JASMINE_*aug*_ represents the augmented JASMINE model with 64 dimensional modality-specific embeddings. For each cancer type, the best result is highlighted in **boldface**. Second best is <u>underlined</u>.

| Dataset/Method | MIND | JASMINE | JASMINE<sub>aug</sub> |
| --- | --- | --- | --- |
| LUAD | **0.275** | 0.076 | <u>0.174</u> |
| KIRC | **0.393** | 0.123 | <u>0.186</u> |
| BRCA | **0.378** | 0.092 | <u>0.169</u> |
| LGG | **0.499** | 0.156 | <u>0.250</u> |
| OV | **0.205** | 0.031 | <u>0.183</u> |
| SKCM | **0.283** | 0.122 | <u>0.189</u> |
| THCA | **0.309** | 0.149 | <u>0.237</u> |
| BLCA | **0.340** | 0.075 | <u>0.212</u> |
| STAD | **0.352** | 0.019 | <u>0.086</u> |
| UCEC | **0.326** | 0.182 | <u>0.236</u> |
| COADREAD | **0.308** | 0.089 | <u>0.168</u> |
| COAD | **0.264** | 0.139 | <u>0.173</u> |
| GBM | **0.203** | 0.119 | <u>0.158</u> |
| HNSC | **0.258** | 0.092 | <u>0.169</u> |
| LUSC | **0.265** | 0.082 | <u>0.176</u> |
| LIHC | **0.417** | <u>0.206</u> | 0.192 |
| CESC | **0.270** | 0.135 | <u>0.199</u> |

In addition to individual datasets, we also combined the 17 TCGA datasets, and apply MIND to the resulting merged dataset consisting of 8,567 patients and 16 cancer types (COAD is a subset of the COADREAD dataset). See Sec 4.4 for details of the merged dataset. Similarly to the individual case, we consider three supervised tasks: cancer type classification, survival prediction and multiomics data reconstruction. For classification, we train XGboost classifiers to predict cancer types using embeddings generated by different methods. For survival prediction and multiomics data reconstruction, we compare the proposed method with IntegraO, JASMINE and MSNE using the same settings as in the individual cases. The survival prediction performance and classification accuracies estimated by 5-fold CV are reported in Table 1, 4 respectively. For each of the five modalities, the Pearson correlations between the predicted and observed are reported in Table 5

**Table 4:** Cancer type classification. Classification accuracy estimated using 5-fold CV. For this task, XGboost classifiers are trained to predict cancer types using embeddings given by different methods. For each cancer type, the best result is highlighted in **boldface**. Second best is <u>underlined</u>.

| Dataset/Method | MIND | IntegraO | JASMINE | MSNE |
| --- | --- | --- | --- | --- |
| TCGA-combined | <u>0.974</u> | 0.948 | **0.975** | 0.891 |
| CCMA | **0.793** | <u>0.761</u> | 0.704 | 0.548 |
| CCLE | **0.659** | 0.547 | <u>0.610</u> | 0.367 |

**Table 5:** Reconstruction. Pearson correlations between prediction and observed values of the individual modalities. JASMINE_*aug*_ represents the augmented JASMINE model with 64 dimensional modality-specific embeddings. For each modality, the best result is highlighted in **boldface**. Second best is <u>underlined</u>.

| Dataset | Modality / Method | MIND | JASMINE | JASMINE<sub>aug</sub> |
| --- | --- | --- | --- | --- |
| TCGA | mRNA | **0.797** | 0.398 | <u>0.554</u> |
|  | DNA methyl | **0.778** | 0.285 | <u>0.398</u> |
|  | CNV | **0.559** | 0.110 | <u>0.149</u> |
|  | RPPA | **0.514** | 0.126 | <u>0.336</u> |
|  | miRNA | **0.741** | 0.358 | <u>0.702</u> |
| CCMA | mRNA | **0.550** | 0.064 | <u>0.106</u> |
|  | DNA methyl | **0.648** | 0.198 | <u>0.431</u> |
|  | CNV | **0.836** | 0.461 | <u>0.548</u> |
| CCLE | RNA | **0.572** | 0.239 | <u>0.280</u> |
|  | DNA methyl | **0.447** | 0.163 | <u>0.219</u> |
|  | CNV | **0.174** | 0.042 | <u>0.072</u> |
|  | miRNA | **0.244** | 0.064 | <u>0.073</u> |
|  | RPPA | **0.379** | 0.108 | <u>0.210</u> |
|  | Metabolomics | **0.338** | 0.121 | <u>0.227</u> |

**Table 6:**
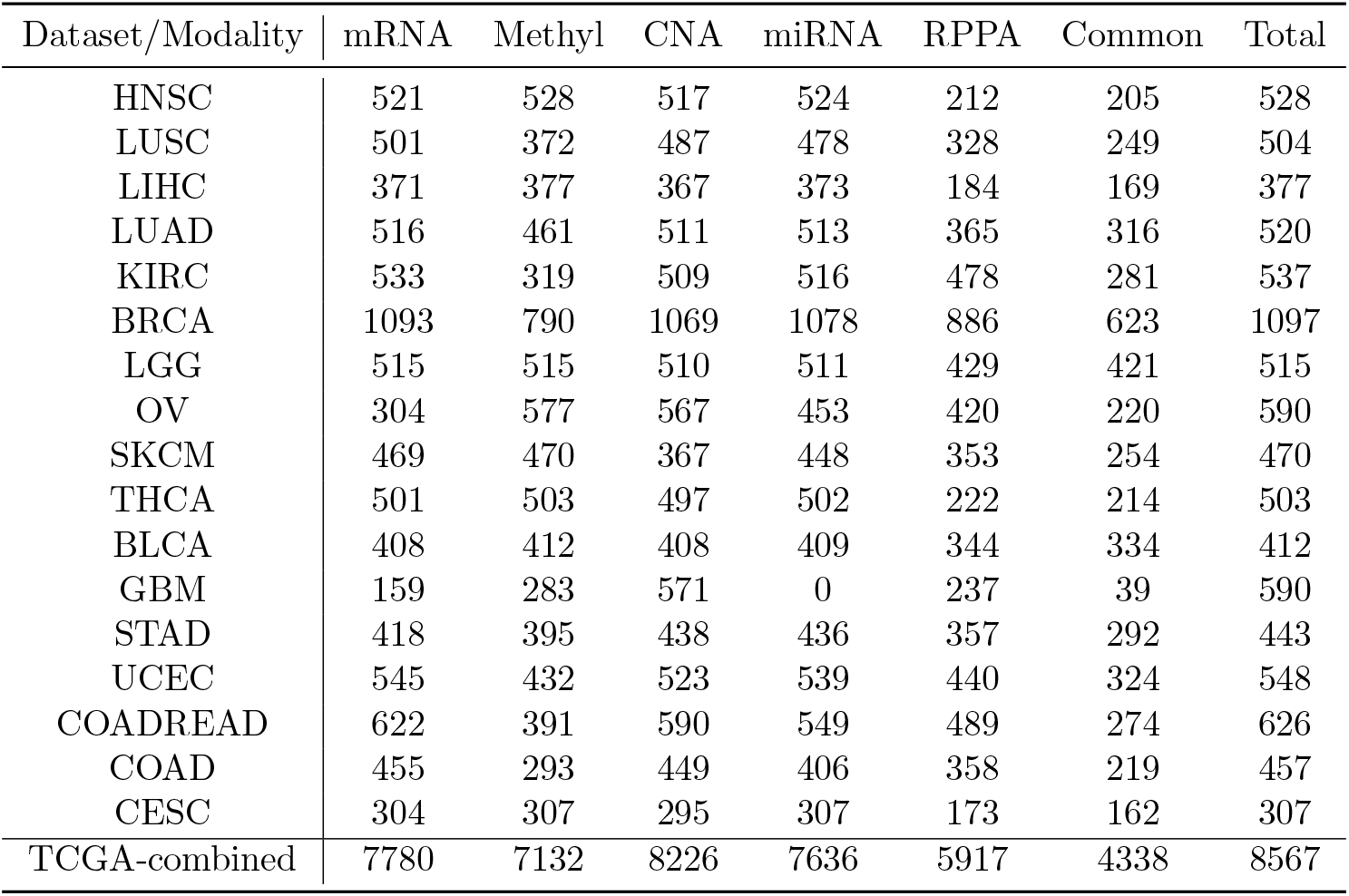
Sample sizes of each dataset investigated in the TCGA example.

| Dataset/Modality | mRNA | Methyl | CNA | miRNA | RPPA | Common | Total |
| --- | --- | --- | --- | --- | --- | --- | --- |
| HNSC | 521 | 528 | 517 | 524 | 212 | 205 | 528 |
| LUSC | 501 | 372 | 487 | 478 | 328 | 249 | 504 |
| LIHC | 371 | 377 | 367 | 373 | 184 | 169 | 377 |
| LUAD | 516 | 461 | 511 | 513 | 365 | 316 | 520 |
| KIRC | 533 | 319 | 509 | 516 | 478 | 281 | 537 |
| BRCA | 1093 | 790 | 1069 | 1078 | 886 | 623 | 1097 |
| LGG | 515 | 515 | 510 | 511 | 429 | 421 | 515 |
| OV | 304 | 577 | 567 | 453 | 420 | 220 | 590 |
| SKCM | 469 | 470 | 367 | 448 | 353 | 254 | 470 |
| THCA | 501 | 503 | 497 | 502 | 222 | 214 | 503 |
| BLCA | 408 | 412 | 408 | 409 | 344 | 334 | 412 |
| GBM | 159 | 283 | 571 | 0 | 237 | 39 | 590 |
| STAD | 418 | 395 | 438 | 436 | 357 | 292 | 443 |
| UCEC | 545 | 432 | 523 | 539 | 440 | 324 | 548 |
| COADREAD | 622 | 391 | 590 | 549 | 489 | 274 | 626 |
| COAD | 455 | 293 | 449 | 406 | 358 | 219 | 457 |
| CESC | 304 | 307 | 295 | 307 | 173 | 162 | 307 |
| TCGA-combined | 7780 | 7132 | 8226 | 7636 | 5917 | 4338 | 8567 |

### 2.4 Childhood Cancer Model Atlas

We applied our proposed method to mRNA-seq, DNA methylation, and copy number variation (CNV) data from the Childhood Cancer Model Atlas (CCMA) dataset (Sun et al., 2023). The dataset consists of multiomics data of 182 paediatric cancer cell lines representing 15 different types of childhood tumour. Details regarding data preprocessing can be found in Supp 4.4. Similarly to the previous examples, we fix the embedding size to 64. We compare the proposed method with IntegrAO, JASMINE and MSNE on cancer type classification and multiomics data reconstruction. The benchmarking strategy is identical to the ones in the previous example. The classification accuracies estimated by 5-fold CV are reported in Table 4. For each of the three modalities, the Pearson correlations between the predicted and observed is reported in Table 5

### 2.5 Cancer Cell Line Encyclopedia

We applied MIND to the Cancer Cell Line Encyclopedia (CCLE) (Ghandi et al., 2019) dataset. The dataset includes RNA-sequencing, DNA methylation, copy number variation (CNV), microRNA, proteomics reverse phase protein array (RPPA), and metabolomics data for 1,461 cells. We benckmark MIND on cancer type classification and data reconstruction following exactly the same procedure as in the previous example. Results are reported in Table 4 and 5 respectively.

## 3 Discussion

Our proposed MIND adopts a multimodal VAE architecture to accommodate incomplete multiomics data. The data-driven prior in MIND injects intrinsic clustering structures present in different modalities into the patient embeddings in an explicit and interpretable fashion, encouraging the learnt embeddings to maximally preserve the clustering information from the observations with arbitrary missing patterns. Compared with current state-of-the-art VAE-based models such as JASMINE (Ballard et al., 2025), another key advantage of MIND is its lightweight, conceptually simple architecture: MIND consists of only 2*M* modality specific encoder and decoder networks, while JASMINE requires training *M* ^2^ + *M* interacting encoders and decoders, making it difficult to scale up. Experiments show that MIND consistently outperforms or achieves performance on par with existing VAE- and network-based state-of-the-art methods on both supervised and unsupervised downstreaming tasks. This suggests that MIND can better extract biologically meaningful information from incomplete multiomics data. Furthermore, we also demonstrate that MIND consistently outperforms JASMINE in terms of predicting unseen data from learnt patient embeddings. This further confirms that MIND is capable of accurately identifying and capturing characteristics of different patients groups in diverse biological contexts.

## Acknowledgements

The authors are supported by an EPSRC Turing AI Acceleration Fellowship (Grant Ref: EP/V023233/1) and the the Pioneer Center for Statistical and computational Methods for Advanced Research to Transform Biomedicine (SMART-Biomed), DNRF grant number P4.

## 4 Methods

Our proposed method can be interpreted as a multimodal variational autoencoder. Before we give the details of the proposed model, we first fix the notations. Let *M* be the total number of modalities. Let *N* be the total number of individual patients. Let [*N* ] = {1, …, *N* }, [*M* ] = {1, …, *M* }. Let 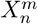 be the data for the *m*th modality of the *n*th patient. For each patient *n*, denote *A*_*n*_ = {*m* : *m* ∈ [*M* ], 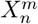 is available} the index set of modalities in which the data of the *n*th patient are available. Similarly, define *B*^*m*^ = {*n* : *n* ∈ [*N* ], 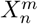 is availabel} be the index set of available patients for each modality. Denote 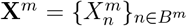 the observed data from the *m*th modality. Denote 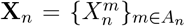 the collection of observed data associated with the *n*th patient. Denote 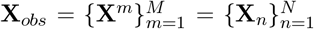 the full multi-omics dataset. In this paper, we assume 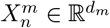 for all *n* ∈ [*N* ], *m* ∈ [*M* ] for sake of clarity. Extension to e.g. count data is straightforward.

We assume that each patient *i* is associated with a latent embedding 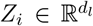 drawn from some unknown distribution. Our goal is to learn the distributions of patient-specific *Z*_*i*_s from the observed **X**_*obs*_. In the following sections, we first discuss the encoder architecture and the choice of prior and posterior distributions on embeddings, then give the training and inference procedure of the proposed model.

### 4.1 Encoder architecture

For each modality *m* ∈ [*M* ], let 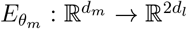 be a modality-specific encoder governed by the parameter vector *θ*_*m*_ that maps 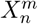, *n* ∈ [*N* ] to the latent space. Denote 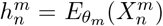 and 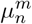,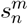 the first and last *d*^*l*^ entries of 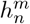, respectively. Denote 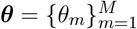 the collection of encoder parameters. Since our goal is to learn patient level embeddings *Z*_*n*_, we need to further aggregate the modality-specific 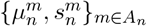 encoded by 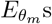, and pass the information to *Z*_*n*_. In this paper, we choose to bridge patient level *Z*_*n*_ and modality-specific 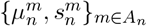 for each patient *n* by setting the VAE posterior

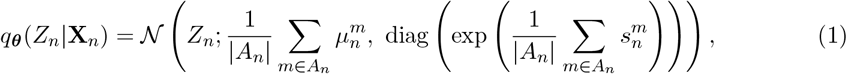

where diag(*z*) represents a diagonal matrix with diagonal elements equal to *z* and off-diagonal elements 0, and exp(*z*) denotes element-wise exponential. In principle, one could consider more sophisticated aggregation functions such as Set Transformer (Lee et al., 2019) instead of simple averaging. However, we found that it gave satisfactory results in all numerical examples. We therefore do not investigate other alternatives in this paper for simplicity.

### 4.2 Prior distribution on embeddings

So far we have discussed encoder architecture and posteriors of the embeddings. In this section, we discuss the choice of prior on *Z*_*n*_s. In this paper, we use a partially data-driven prior on *Z*_*n*_s to inject the neighbouring structure information into the embeddings. Our choice of prior is inspired by the variational formulation of t-SNE (Maaten and Hinton, 2008) given in Ravuri et al. (2023). 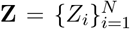 and 𝒟 (**Z**) ∈ ℝ ^*N×N*^ be the pairwise similarity matrix of embeddings **Z** where each entry 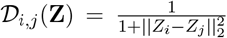. For each modality *m* ∈ [*M* ], let 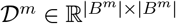 be the t-SNE’s sparse data affinity matrix obtained by applying t-SNE to **X**^*m*^. Let 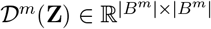 be a sub-matrix of 𝒟 (**Z**) whose rows and columns are selected to match 𝒟^*m*^. Let 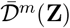, 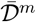 be the normalised and vectorised versions of 𝒟^*m*^(**Z**), 𝒟^*m*^, respectively. Denote *I*_*p*_ a *p* × *p* identity matrix. We define the exponential-tilted prior on **Z** as

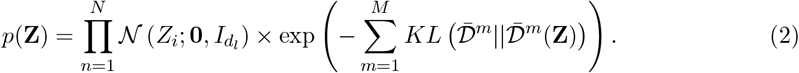

Note that 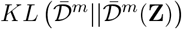, the KL divergence between two categorical distributions specified by probability vectors 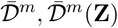, respectively, is nonnegative by definition. This ensures that *p*(**Z**) is still a valid probability distribution whose probability density is known up to a multiplicative constant. Compared to i.i.d. isotropic Gaussian priors, our choice of *p*(**Z**) additionally encourages different subsets of **Z** to cluster in a way similar to the neighbouring structures of **X**^*m*^s, which are encoded as t-SNE’s data affinity matrices. This additional information helps the patient level embeddings **Z** better capture the cluster structure from **X**_*obs*_, and promote information sharing in the latent space. In this paper, the affinity matrices 𝒟 ^*m*^s are computed using t-SNE with the perplexity parameter set to 30, as this choice gives satisfactory results in all numerical examples.

### 4.3 Training and inference

We discuss here the training and inference procedures of the proposed model. Let 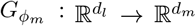 be the modality-specific decoder governed by the parameter vector *ϕ*_*m*_ for each *m* ∈ [*M* ]. Let 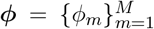 be the collection of decoder parameters. The training procedure of the proposed model is similar to a VAE (Kingma and Welling, 2014): Denote 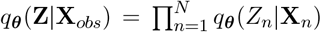 the joint posterior distribution of **Z**. Suppose 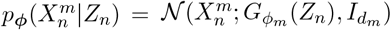, the evidence lower bound of the proposed model takes the form

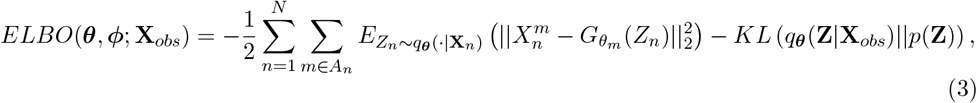

where 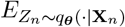 means taking expectation w.r.t. *Z*_*n*_ ∼ *q*(_***θ***_·|**X**_*n*_). The first and second term of *ELBO*(***θ, ϕ***; **X**_*obs*_) are the log likelihood and the KL divergence between the posterior and prior, respectively. Here we assume 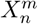 follows a Gaussian distribution. Extension to other likelihoods is straightforward.

In addition to the standard ELBO of a VAE model, we also incorporate a regularisation term on the encoder outputs 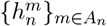 for all *n* ∈ [*N* ] taking the form

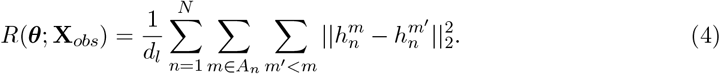

Intuitively speaking, *R*(***θ***; **X**_*obs*_) penalises pairwise distance between 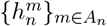 for each pa-tient *n*. Recall that 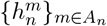 are generated using different modality-specific encoders. We encourage 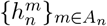 to be close to each other as the data used to generate these quantities are from the same patient, and we want the modality-specific encoders 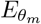 to be aware of this cross-modality connection.

The resulting loss function of the proposed model is

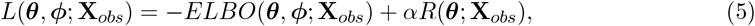

where *α >* 0 is a hyperparameter controlling the strength of the regulariser *R*(***θ***; **X**_*obs*_). We set *α* = 0.05 in all numerical examples. The VAE parameters {***θ, ϕ***} are trained using reparameterisation trick (Kingma and Welling, 2014) and gradient descent methods. Note that the prior we proposed in Sec 4.2 does not factorise. As a result, we are not able to directly apply standard minibatch stochastic gradient descent. We use a Gibbs sampling style training scheme to address this issue: Denote *BC* the index set of a minibatch of patients. Denote **Z**_*BC*_ the corresponding embeddings and **Z**_−*BC*_ = **Z** \ **Z**_*BC*_ its complement. For each minibatch stochastic gradient descent step, we replace *KL* (*q*_***θ***_(**Z**|**X**_*obs*_)||*p*(**Z**)) in Eq (3) by the conditional version *KL* (*q*_***θ***_(**Z**_*BC*_|**Z**_−*BC*_, **X**_*obs*_)||*p*(**Z**_*BC*_|**Z**_−*BC*_)). Since the posterior *q*(**Z**|**X**_*obs*_) is factorisable by design, we have

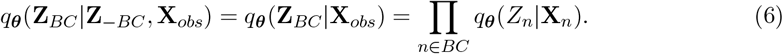

The second term *p*(**Z**_*BC*_|**Z**_−*BC*_) is equivalent to *p*({**Z**_*BC*_, sg(**Z**_−*BC*_)} in the training process, where sg is the stop-gradient operator (i.e. we treat **Z**_−*BC*_ as fixed by detaching it from the computation graph). Computing the corresponding minibatch log likelihood and regularisation is straightforward. The final minibatch loss is obtained by combing the individual minibatch terms the same way as in Eq (5).

Suppose 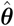,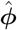 are approximate minimisers of *L*(***θ, ϕ***; **X**_*obs*_). Then each patient embedding *Z*_*n*_ is represented by the posterior 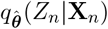 We can also compute the reconstructed or predicted data 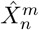 for any *n* ∈ [*N* ], *m* ∈ [*M* ] by passing either the mean or a sample from 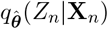 to the corresponding trained decoder 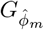. See Fig 1 for a schematic illustration of the proposed method.

## Appendix

### 4.4 Data preprocessing

#### 4.4.1 Synthetic data

Synthetic data are generated using the `InterSim`CRAN package (Chalise et al., 2016). We follow Ma et al. (2025), setting the number of patient *N* = 500 and number of groups *C* = 15. The low noise data are generated using parameters `effect`=1.5 and `p.DMP`=0.1. The high noise data are generated using parameters `effect`=1 and `p.DMP`=0.1.

#### 4.4.2 TCGA

For the TCGA example, we leveraged TCGA multi-omics data sets across 17 types of cancer: Head and Neck squamous cell carcinoma (HNSC), Lung squamous cell carcinoma (LUSC), Liver hepatocellular carcinoma (LIHC), Cervical and endocervical cancers (CESC), Lung adenocarcinoma (LUAD), Kidney renal clear cell carcinoma (KIRC), Breast invasive carcinoma (BRCA), Brain Lower Grade Glioma (LGG), Ovarian serous cystadenocarcinoma (OV), Skin Cutaneous Melanoma (SKCM), Thyroid carcinoma (THCA), Bladder urothelial carcinoma (BLCA), Glioblastoma multiforme (GBM), Stomach adenocarcinoma (STAD), Uterine Corpus Endometrial Carcinoma (UCEC), Colorectal adenocarcinoma (COADREAD), and Colon adenocarcinoma (COAD). We obtain mRNA expression, DNA methylation, MicroRNA, protein expression data and relavent patient information from the [Broad Institute’s Firehose source data](https://gdac.broadinstitute.org/). Copy number variation are obtained from [cBioportal](https://www.cbioportal.org/). For the merged data, we first combine the 17 data sets, then remove columns with more than 20% missing values for each of the modalities. We then impute missing values in the resulting datasets using *k*-nearest neighbour. For copy number variation, mRNA expression, and DNA methylation data, we select the top 2,000 most variable features. Data from all modalities are then standardised so that each feature dimension has mean 0 and standard deviation 1. Datasets for each individual cancer type are preprocessed in the same fashion.

#### 4.4.3 CCMA

The CCMA dataset consists of three modalities: RNA-sequencing, DNA methylation, and copy number variation. The CCMA dataset and relevant patient information are obtained from the [web portal](https://data.mendeley.com/datasets/rnfs539pfw/3). Missing values are imputed using *k*-nearest neighbour. For RNA-sequence and DNA methylation data, we select the top 2000 most variable features. All data are then standardised so that each feature dimension has mean 0 and standard deviation 1.

#### 4.4.4 CCLE

The CCLE dataset consists of six modalities: RNA-sequencing, DNA methylation, copy number variation, MicroRNA, protein expression data and metabolomics data. The CCLE dataset and relevant patient information are obtained from the [web portal](https://depmap.org/portal/). For RNA-sequence and DNA methylation data, we select the top 2000 most variable features. All data are then standardised so that each feature dimension has mean 0 and standard deviation 1.

### 4.5 Data availability and reproducibility

Implementation of the proposed method, as well as codes for reproducing all experiments in the paper, can be found on the [GitHub page](https://github.com/hwxing3259/MIND). All datasets and output files can be downloaded from [Figshare](https://figshare.com/articles/dataset/Multi_O_Int/30032023).

